# Hexokinase 2 displacement from mitochondria-associated membranes prompts Ca^2+^-dependent death of cancer cells

**DOI:** 10.1101/736538

**Authors:** Francesco Ciscato, Riccardo Filadi, Ionica Masgras, Marco Pizzi, Oriano Marin, Nunzio Damiano, Paola Pizzo, Alessandro Gori, Federica Frezzato, Livio Trentin, Paolo Bernardi, Andrea Rasola

## Abstract

Cancer cells undergo changes in metabolic and survival pathways that increase their malignancy. Isoform 2 of the glycolytic enzyme hexokinase (HK2) enhances both glucose metabolism and resistance to death stimuli in many neoplastic cell types. Here we observe that HK2 locates at mitochondria-endoplasmic reticulum (ER) contact sites called MAMs (Mitochondria-Associated Membranes). HK2 displacement from MAMs with a selective peptide triggers mitochondrial Ca^2+^ overload caused by Ca^2+^ release from ER via inositol-3-phosphate receptors (IP3Rs) and by Ca^2+^ entry through plasma membrane. This results in Ca^2+^-dependent calpain activation, mitochondrial depolarization and cell death. The HK2-targeting peptide causes massive death of chronic lymphocytic leukemia B cells freshly isolated from patients, and an actionable form of the peptide reduces growth of breast and colon cancer cells allografted in mice without noxious effects on healthy tissues. These results identify a signalling pathway primed by HK2 displacement from MAMs that can be activated as anti-neoplastic strategy.

## Introduction

Hexokinases are a family of four isoforms that catalyze phosphorylation of glucose, making it available for utilization in glycolysis, pentose phosphate pathway, glycogenesis and hexosamine biosynthesis (Wilson, 2003). HK2, the most active isozyme, is markedly expressed in cells characterized by a high rate of glucose consumption, such as adipose, skeletal and cardiac muscle. During the neoplastic process, metabolic changes are required to allow cell growth in conditions of fluctuating nutrient and oxygen availability (Vander Heiden & DeBerardinis, 2017). HK2 plays a major role in this metabolic rewiring (DeWaal *et al*, 2018; Robey & Hay, 2006; Wang *et al*, 2014), being induced by oncogenic K-Ras activation (Patra *et al*, 2013) or in response to (pseudo)hypoxia (Bhalla *et al*, 2018; Semenza, 2013). HK2 is mainly bound to the outer mitochondrial membrane, where it can gain privileged access to newly synthesized ATP, thus increasing efficiency in glucose usage (Mathupala *et al*, 2009), while following glucose deprivation HK2 elicits autophagy by inhibiting mTORC1 (Roberts *et al*, 2014). HK2 binding to mitochondria is increased by Akt phosphorylation, a key event occurring downstream to many signalling pathways hyperactivated in tumor cells (Miyamoto *et al*, 2008). Moreover, mitochondrial HK2 takes part in the protection of cancer cells from noxious stimuli through poorly defined mechanisms that include antagonizing the activity of pro-apoptotic Bcl-2 family proteins and increasing anti-oxidant defenses through pentose phosphate pathway induction (Roberts & Miyamoto, 2015). In cancer patients, HK2 induction is related to stage progression, acquisition of invasive and metastatic capabilities and poor prognosis (Mathupala *et al*, 2010). HK2 promotes neoplastic growth in glioblastoma multiforme (Wolf *et al*, 2011), confers chemoresistance in epithelial ovarian cancer (Suh *et al*, 2014) and is required for tumor onset and maintenance in mouse models of lung and breast cancer, where its genetic ablation is therapeutic without adverse effects (Patra et al., 2013).

Thus, HK2 constitutes a promising target for developing anti-neoplastic strategies, but the clinical use of hexokinase inhibitors is hampered by lack of specificity or side effects (Akins *et al*, 2018) potentially associated with glucose metabolism derangement. A possible alternative approach is detaching HK2 from mitochondria, as we and others have previously shown that this can induce opening of a mitochondrial channel, the permeability transition pore (PTP), and consequently cell death (Chiara *et al*, 2008; Masgras *et al*, 2012; Pantic *et al*, 2013; Roberts & Miyamoto, 2015). However, both a detailed comprehension of the molecular mechanisms leading to cell damage and the development of a HK2-targeting tool that is operational in *in vivo* tumor models are required to translate this information into the groundwork for future anti-neoplastic approaches.

Here we demonstrate that in neoplastic cells HK2 localizes in MAMs, specific subdomains of interaction between mitochondria and ER. HK2 detachment from MAMs rapidly elicits a massive Ca^2+^ flux into mitochondria and consequently a calpain-dependent cell death. We ignite this process with a HK2-targeting peptide composed by modular units that can be adapted to *in vivo* delivery, without affecting hexokinase enzymatic activity and with no adverse effects on animal models.

## Results

### HK2 localizes in MAMs of neoplastic cells

Dissection of sub-mitochondrial HK2 localization can provide important functional clues, as mitochondria compartmentalize specific activities in domains formed by multiprotein platforms. After confirming that HK2 is expressed in diverse tumor samples (Fig 1A and 1B) and associates with mitochondria (Fig 1C), we have found that HK2 specifically localizes in MAMs by merging the fluorescence of HK2-conjugated antibodies with mitochondria-targeted YFP and ER-targeted CFP (Fig 1D), or with a split-GFP-based probe for ER-mitochondria contacts (SPLICS_L_) (Cieri *et al*, 2018) (Fig 1E-1G and Fig EV1A). This same analysis showed that HK2 is significantly enriched in MAMs with respect to TOM20, a protein that is uniformly distributed in the outer mitochondrial membrane (Fig EV1B). MAMs are dynamic structures that control the exchange between ER and mitochondria of ions and lipids, tuning complex biological processes such as ER stress, autophagy, cell death and maintenance of glucose homeostasis (Csordas *et al*, 2018; Prudent & McBride, 2017; Rieusset, 2018). A pivotal role of MAMs is the regulation of Ca^2+^ fluxes from ER to mitochondria through IP_3_Rs (Filadi *et al*, 2017); thus, HK2 displacement from MAMs could affect intracellular Ca^2+^ dynamics, raising the possibility that a Ca^2+^ dyshomeostasis can ensue and damage neoplastic cells.

**Figure 1.**
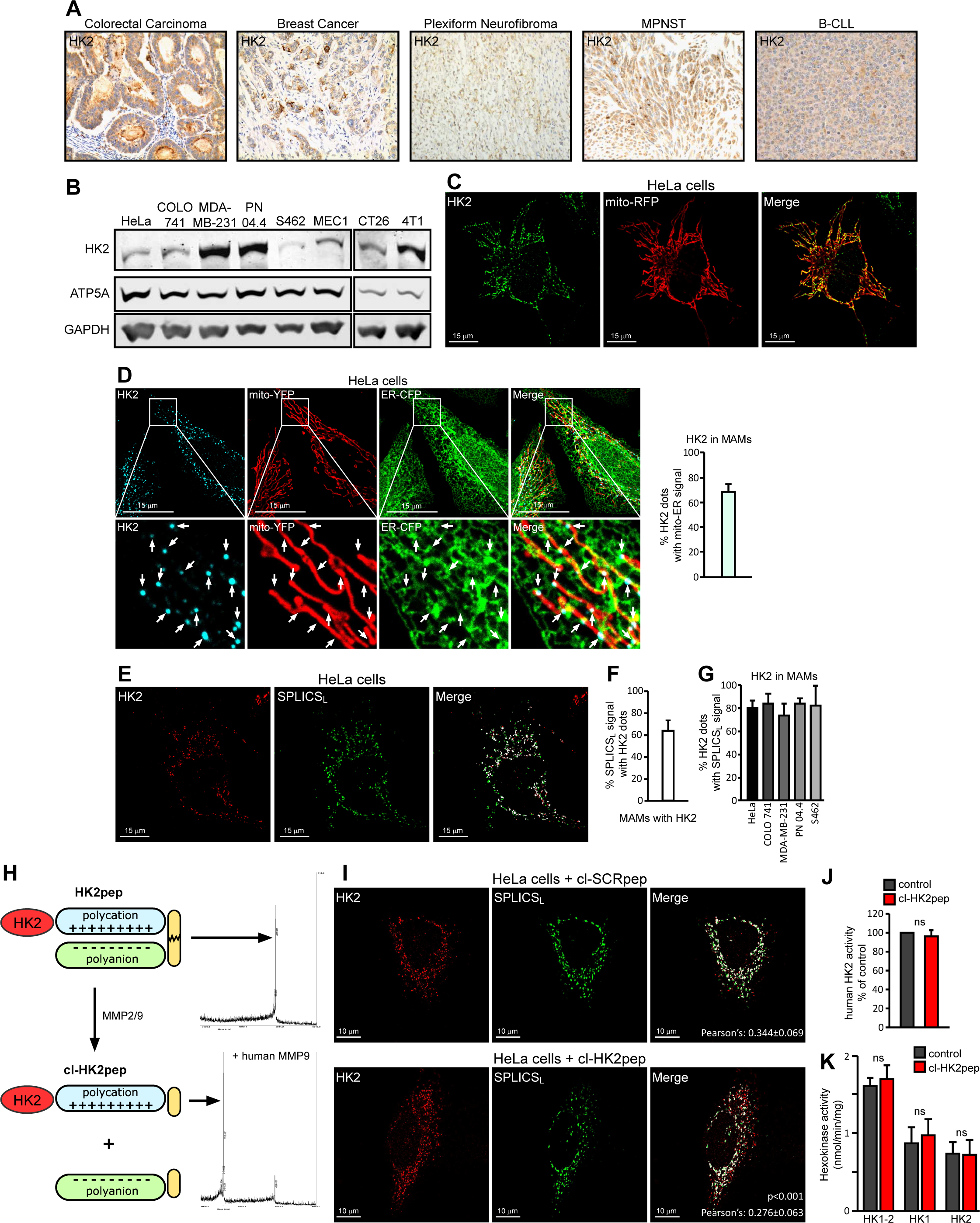
HK2 locates in MAM of cancer cells and is displaced by HK2pep. A, B Analysis of HK2 expression in human and mouse cancer samples by immunohistochemistry (A) and Western immunoblot analysis (B) in neoplastic cell models of human (cervix carcinoma HeLa cells, colorectal carcinoma COLO 741 cells, breast cancer MDA-MB-231 cells, plexiform neurofibroma PN 04.4 cells, malignant peripheral nerve sheath tumor S462 cells, B-chronic lymphocytic leukaemia MEC1 cells) and mouse (colon carcinoma CT26 cells and breast cancer 4T1 cells) origin. ATP5A and GAPDH are mitochondrial and cytosolic loading controls, respectively. C Immunofluorescence staining of HK2 with an AlexaFluor488-conjugated antibody in HeLa cells expressing mitochondria-targeted RFP. Yellow signals in the merge analysis indicate mitochondrial localization of HK2. Scale bar: 15µM. D Immunofluorescence staining of HK2 with a secondary AlexaFluor555-conjugated antibody in HeLa cells expressing both mitochondria-targeted YFP and ER-targeted CFP. The merged white signal indicates MAM localization of HK2 and is quantified in the bar graph on the right (n=24). Image magnifications are shown in the lower part of the panel; arrows indicate HK2 dots in mito-ER contact sites. Scale bar: 15µM. E-G Fluorescence co-staining of HK2 and split-GFP-based probe for ER-mitochondria contacts (SPLICS_L_) on HeLa cells; HK2 is revealed with a secondary AlexaFluor555-conjugated antibody, the merged signal is white (E; scale bar: 15µM); (F) percentage of SPLICS_L_ dots positive for HK2 (n=10 cells); (G) percentage of HK2 dots positive for SPLICS_L_ (n≥6 cells). H Functional unit composition of HK2pep (left); the HK2-targeting sequence is in red; the polycation and polyanion stretches are in light blue and light green, respectively; the MMP2/9 target sequence is in yellow. On the right, mass spectrometry profile of HK2pep before and after incubation (cl-HK2pep) with human MMP9. I HK2 displacement from HeLa MAMs after a 2 min treatment with cl-HK2pep is shown by loss of merging signal analyzed as in (E). cl-SCRpep is used as a negative control and Pearson’s co-localization coefficient is indicated in figure (cl-SCRpep n=26 cells; cl-HK2pep n=24 cells; p<0.001 with a Student’s *t* test). Scale bar: 10µM. J, K Effect of cl-HK2pep on glucose phosphorylation by human recombinant HK2 (J) or in 4T1 cell extracts (K), where both total hexokinase activity and HK1/HK2 specific activities are measured.

### Design of a peptide that displaces HK2 form MAMs without affecting hexokinase enzymatic activity

To investigate this possibility and to generate a tool with a potential anti-tumor activity *in vivo*, we have conceived a HK2-targeting Cell Penetrating Peptide (CPP), dubbed HK2pep (Fig 1H). HK2pep is designed with a modular structure composed of: i) a N-terminal HK2 tail, which acts as the active moiety by displacing HK2 from the outer mitochondrial membrane (in the negative control, SCRpep, this HK2-specific sequence is substituted by a scrambled one); ii) a polycation stretch required for plasma membrane permeation; iii) a polyanion sequence that shields polycation charges; iv) a matrix metalloprotease 2 and 9 (MMP2/9) target sequence that links the two charged stretches. As previously observed with similar actionable CPPs (Olson *et al*, 2009), the metalloprotease target sequence inhibits cell uptake of the peptide until its polycation sequence is unmasked by MMP2/9 cleavage. MMPs are highly expressed in a variety of tumor types, where they induce extracellular matrix remodeling and favor cancer cell invasiveness (Isaacson *et al*, 2017). Hence, HK2pep should be preferentially activated inside neoplasms, and its subsequent entry through the plasma membrane would then lead to HK2 displacement from mitochondria and eventually cancer cell death. Moreover, HK2pep is not permeable across the endothelium of normal blood vessels, but its dimension (about 5 kDa) allows passage across the fenestrated endothelium that perfuse many cancer types.

The active moiety of HK2pep (*i.e.* the cleaved peptide, cl-HK2pep, Fig 1H) is indeed able to enter cells, to interact with mitochondria (Fig EV2A) and to induce HK2 redistribution from MAMs into cytosol in less than 2 minutes (Fig 1I and Fig EV2B). cl-HK2pep does not perturb hexokinase enzymatic activity either on the purified enzyme (Fig 1J) or on cell samples, where it is equally ineffective on HK2 and on the widespread isozyme HK1 (Fig 1K and Fig EV2C).

### HK2 detachment from MAMs elicits a Ca^2+^ flux into mitochondria via IP3Rs and plasma membrane that causes mitochondrial depolarization

In accord with a role played by several MAM proteins in the regulation of Ca^2+^ homeostasis, cl-HK2pep prompts cycles of ER Ca^2+^ release and refill (Fig 2A) and boosts cytosolic IP_3_ levels (Fig 2B). This IP_3_ rise is prevented by pre-incubation with the IP_3_R inhibitor Xestospongin-C (Xe-C; Fig 2B) and by chelating cytosolic Ca^2+^ (Fig EV3A), and delayed with respect to the Ca^2+^ release from ER (compare Fig 2A and 2B). These observations are consistent with cl-HK2pep eliciting a primary Ca^2+^ efflux from ER that prompts a surge in cytosolic IP_3_ (Rebecchi & Pentyala, 2000), further amplifying ER Ca^2+^ release via IP_3_Rs.

**Figure 2.**
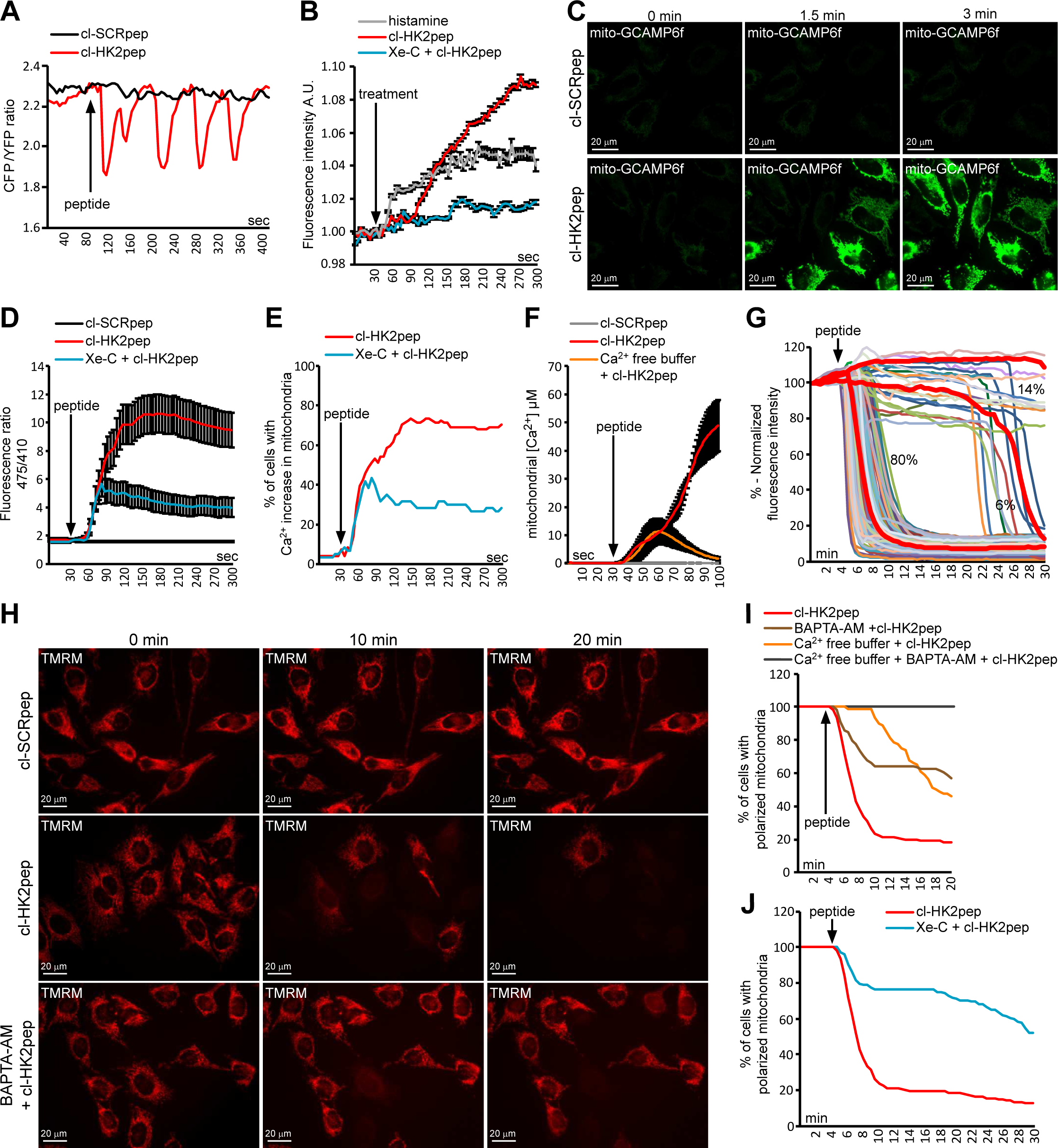
HK2 detachment from MAMs prompts Ca^2+^ influx and depolarization in mitochondria following IP3R opening. A-F Effect of cl-HK2pep on cellular Ca2^+^ dynamics and IP_3_ levels. ER Ca^2+^ levels are measured by the FRET-based, D4ER fluorescent probe expressed in the lumen of ER (A); IP_3_ levels are assessed with the GFP-PHD probe; histamine (100μM) is used as a positive control for IP_3_ generation; data are reported as mean of fluorescent signals±SEM (n=3; B); changes in mitochondrial Ca^2+^ levels are recorded (C; scale bar: 20µM) and quantified using the GCAMP6f sensor (in D, as mean of 475/410 nm ratio signal±SEM, in (E) as percentage of cells with increased Ca^2+^ in mitochondria, with a threshold 475/410 ratio for positivity >3; baseline mean ratio=1.84±0.54) or with mitochondria-targeted aequorin (F, where data are reported as mean of [Ca^2+^]±SEM). G-J Effect of cl-HK2pep treatment on mitochondrial membrane potential assessed with the TMRM probe. Kinetic experiments (G, single cell analysis, n=216; H, representative field; scale bar: 20µM) are quantified (I and J; TMRM fluorescence is normalized to initial value and expressed in percentage for each time point, with a depolarization threshold placed at 40% of initial value). Experiments throughout the Figure are carried out on HeLa cells; cl-SCRpep, negative control of cl-HK2pep (2 μM each). Where indicated, cells are kept in Ca^2+^ free medium plus 500 µM EGTA with or without 10 µM BAPTA-AM; Xe-C is the IP3R inhibitor Xestospongin C (5 μM).

Mitochondria can promptly take up Ca^2+^ released from ER, resulting in modulation of the activity of Krebs cycle enzymes and preventing deregulated increases in cytosolic [Ca^2+^] (Cannino *et al*, 2018; Filadi et al., 2017). cl-HK2pep treatment rapidly boosts mitochondrial [Ca^2+^] in a stable and Xe-C-sensitive way (Movie 1 and Figure 2C-2E), rising mitochondrial [Ca^2+^] to about 50 µM (Fig 2F). However, chelation of extracellular Ca^2+^ only induces a transient mitochondrial [Ca^2+^] peak of about 12 µM (Fig 2F), indicating that cl-HK2pep administration elicits both Ca^2+^ release from ER and Ca^2+^ entry through plasma membrane (Berridge *et al*, 2003) and that mitochondria take up Ca^2+^ from both sources. HK2 targeting also prompts a sudden and massive mitochondrial depolarization (Fig 2G and 2H, Movie 2 and Fig EV3B-EV3E), which follows the increase in mitochondrial [Ca^2+^] (Fig EV3F and EV3G). This depolarization is inhibited by chelating cytosol or extracellular Ca^2+^ and by the IP_3_R inhibitor Xe-C (Fig 2I and 2J, Movie 3 and Fig EV3H). Therefore, the HK2-targeted peptide causes mitochondrial Ca^2+^ overload as a consequence of Ca^2+^ release form ER via IP_3_Rs and of Ca^2+^ entry through plasma membrane.

### The HK2-targeting peptide triggers calpain-dependent cell death

Overcoming the efflux and the buffering capacity of Ca^2+^ in mitochondria can induce the permeability transition pore (PTP), a high conductance channel the opening of which commits cells to death (Rasola & Bernardi, 2011; Rasola & Bernardi, 2014). PTP opening is independently inhibited by two unrelated molecules, Cyclosporin-A (CsA) and C63, but this inhibitory effect can be overwhelmed by intense stimuli of PTP induction (Bernardi *et al*, 2015). We find that both CsA and C63 do not affect mitochondrial Ca^2+^ uptake following cl-HK2pep treatment (Fig 3A), but markedly delay mitochondrial depolarization (Fig 3B), indicating that this occurs downstream to PTP induction.

**Figure 3.**
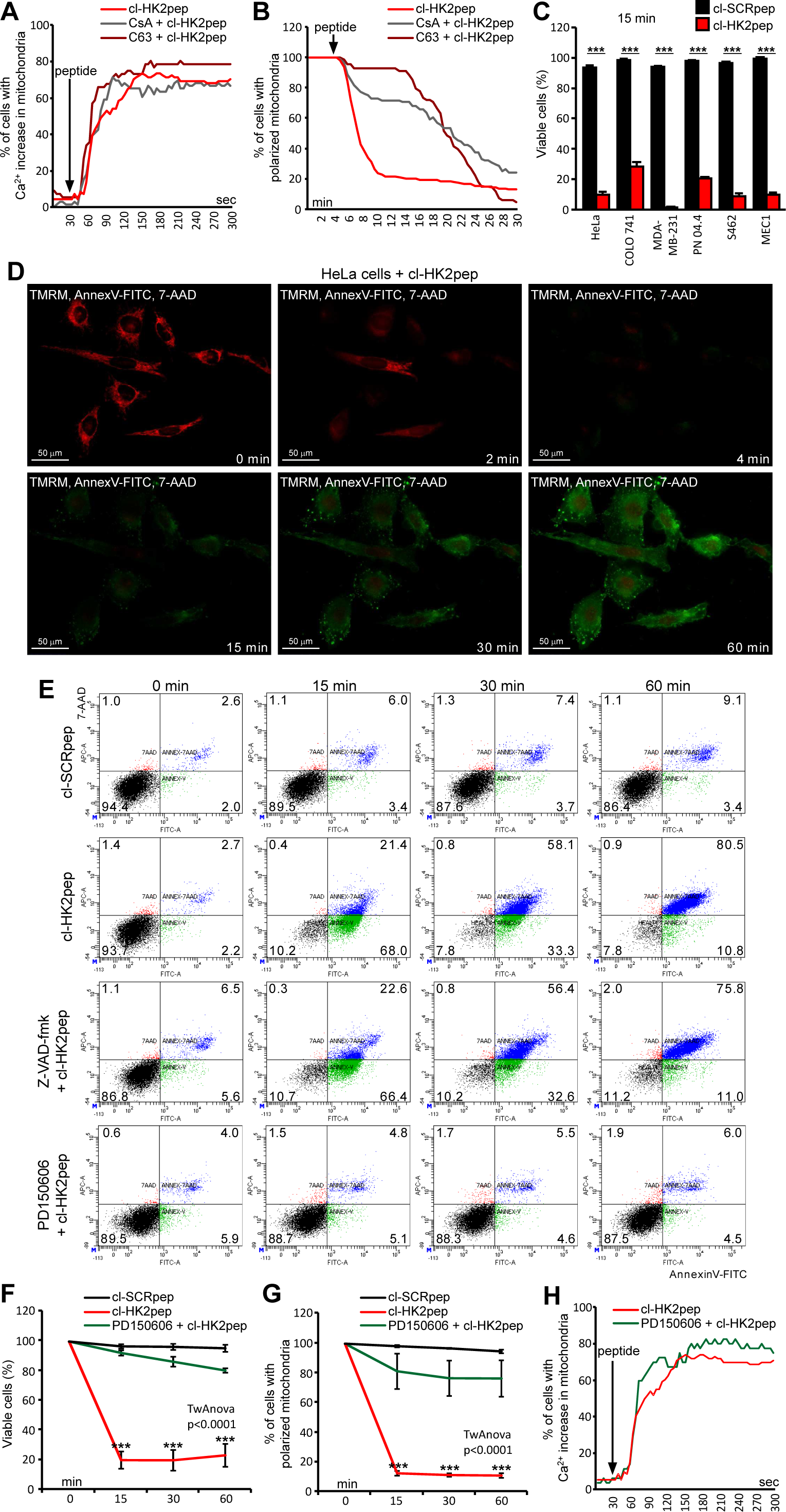
HK2pep induces PTP- and calpain-dependent cell death. A, B Effects of the PTP desensitizers CsA or C63 (5 µM each, 1 h preincubation) on mitochondrial Ca^2+^ levels recorded with GCAMP6f (A) and on mitochondrial membrane potential assessed with TMRM (B) in cells treated with cl-HK2pep. C-F cell death induction by cl-HK2pep; in the cytofluorimetric analyses reported in (C, E and F), viable cells are double negative for Annexin V-FITC and 7-AAD staining and measured 15 min after peptide treatment; in the kinetic experiment shown in (D; scale bar: 50µM) viable cells are double negative for Annexin V-FITC and 7-AAD and TMRM positive. G, H Effect of calpain inhibition on mitochondrial membrane potential assessed with TMRM (G) and on mitochondrial Ca^2+^ levels recorded with GCAMP6f (H) in cells treated with cl-HK2pep. Experiments throughout the Figure are carried out on HeLa cells; cl-SCRpep, negative control of cl-HK2pep (2 μM each); where indicated, the caspase inhibitor Z-VAD-fmk or the calpain inhibitor PD150606 (50 µM each) are pre-incubated 1 h before peptide treatment. Experiments using TMRM or GCAMP6f probes are analyzed as in Figure 2. In (C) *** p<0.001 with a Student’s *t* test; in (F, G), p<0.0001 with a two-way ANOVA; Bonferroni post-test in graph ***p<0.001.

The HK2-targeting peptide abruptly elicits cell death in all tested cancer cell models (Fig 3C), whereas it is poorly effective in non-transformed cell types expressing HK2 (Fig EV4A-EV4C). Peptide administration triggers mitochondrial depolarization in most tumor cells after 2-4 minutes, followed by phosphatidylserine exposure on the cell surface and plasma membrane rupture in less than one hour (Fig 3D, 3E and Movie 4). Even though these are typical apoptotic changes, none of them is affected by the pan-caspase inhibitor Z-VAD-fmk (Fig 3E and Fig EV4D-EV4F), However, the broad spectrum calpain inhibitor PD150606 abrogates cl-HK2pep-dependent induction of mitochondrial depolarization and cell death (Fig 3E-G and Fig EV4G-EV4M) without affecting the mitochondrial Ca^2+^ rise triggered by the peptide (Fig 3H). Taken together, these data indicate that HK2 displacement from MAMs does not trigger a classical apoptotic pathway, but rather activates a cell death process relying on the Ca^2+^-dependent proteases calpains (Storr *et al*, 2011).

### The HK2-targeting peptide kills primary B-Chronic Lymphatic Leukemia cells and inhibits neoplastic growth of colon and breast cancer cells

In order to test its efficacy as a potential anti-neoplastic treatment, we studied the effect of cl-HK2pep on freshly isolated leukemia cells obtained from B-Chronic Lymphatic Leukemia (B-CLL) patients. B-CLL is the most common leukaemia form that accounts for about 40% of all adult leukemias (Fabbri & Dalla-Favera, 2016). Although in the last years several new molecules, including anti-CD20 antibodies as well as kinase inhibitors, have become available for B-CLL therapy, the disease remains incurable and B-CLL patients usually relapse and/or become refractory. Induction of HK2 contributes to rituximab resistance (Gu *et al*, 2018), and release of MMP9 increases motility of B-CLL cells (Martini *et al*, 2017); hence, B-CLL constitutes an interesting model for testing the effectiveness of HK2pep. All B-CLL patient cells that we have analyzed express HK2 (Fig 4A), independently of their mutation status (Table 1), and undergo a massive mitochondrial depolarization and cell death in the first 15 minutes of cl-HK2pep treatment (Fig 4B-4D). These events are calpain-dependent (Fig 4B-4D), and in all analyzed samples HK2 targeting completely erases B-CLL cells (Fig 4E), whereas it is less effective in inducing death in non-neoplastic, CD19^+^ B cells (Fig 4F).

**Table 1.** Biological and clinical characteristics of the 22 B-CLL patients analyzed. *Mutated was defined as having a frequency of mutations greater than 2% from the germline IGHV sequence. ^§^N = normal karyotype. WBC, white blood cell.

|  |  |
| --- | --- |
| Median age, years (range) | 74 (62-86) |
| Male/Female | 12/10 |
| WBC count, /mm<sup>3</sup> (range) | 72,435 (21,410-168,940) |
| Lymphocytes, % (range) | 80 (58-92) |
| Mutated* | 14/22 |
| Karyotype (N<sup>§</sup>/13q-/12+/11q-/17p-) | 3/15/3/0/1 |

**Figure 4.**
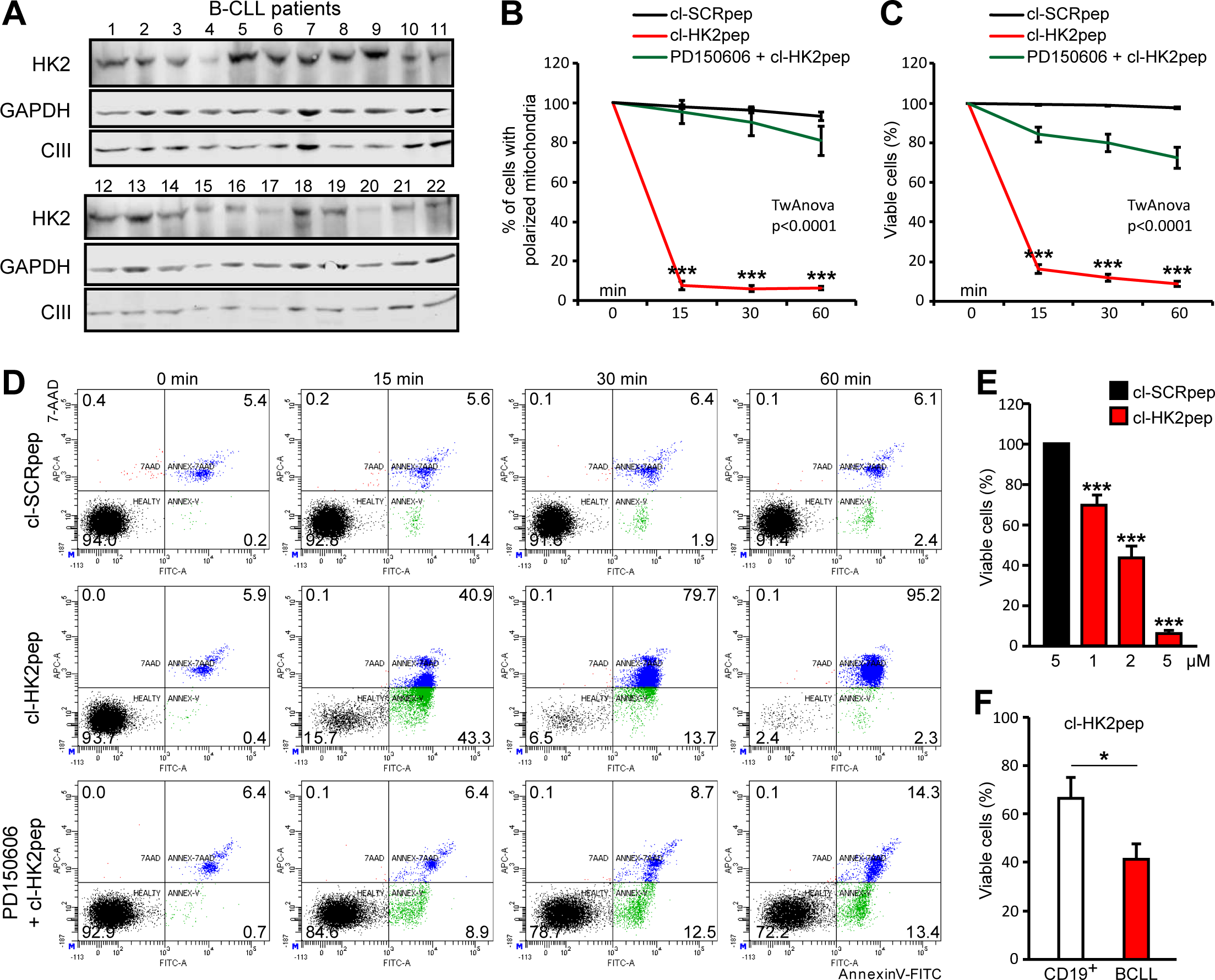
HK2pep kills primary B-CLL cells. A-F Effect of cl-HK2pep on human B-CLL cells freshly isolated from patients. HK2-expressing B-CLL cells (A; GAPDH and respiratory complex III subunit UQCRC2 are cytosol and mitochondrial loading controls, respectively) undergo mitochondrial depolarization assessed by TMRM staining (B) and cell death, measured by cytofluorimetric analysis of Annexin V-FITC and 7-AAD staining (C-F). Cells are treated with 5 µM cl-HK2pep; PD150606 (50 μM) is pre-incubated for 1 h. Experiments are carried out on samples from at least 10 patients and on CD19^+^, primary B lymphocytes from 5 healthy controls. In (B, C) two-way ANOVA: p<0.0001; Bonferroni post-test in graph ***p<0.001, in (E, F), Student’s *t* test; ***p<0.01, *p<0.05.

We have then evaluated the effect of targeting HK2 on solid tumors. Silencing HK2 expression hampers the ability of cancer cells to form colonies (Fig 5A and 5B), and HK2 targeting with both cl-HK2pep and the entire HK2pep inhibits *in vitro* tumorigenic growth by killing cancer cells (Fig 5C and 5D). Efficacy of the entire HK2pep indicates that its active moiety is released by MMP2/9 cleavage, and that this peptide can be used on *in vivo* neoplastic models in which HK2 and MMP2/9 are expressed (Fig 5E and Fig EV5A). We observe that intra-tumor injections of either cl-HK2pep or entire HK2pep significantly decrease the volume of allograft-injected colon cancer cells (Fig 5F), and the same result is achieved by intraperitoneal injection of entire HK2pep on both colon and breast cancer allografts (Fig 5H and 5I), without any tissue damage in treated animals (Fig EV5B).

**Figure 5.**
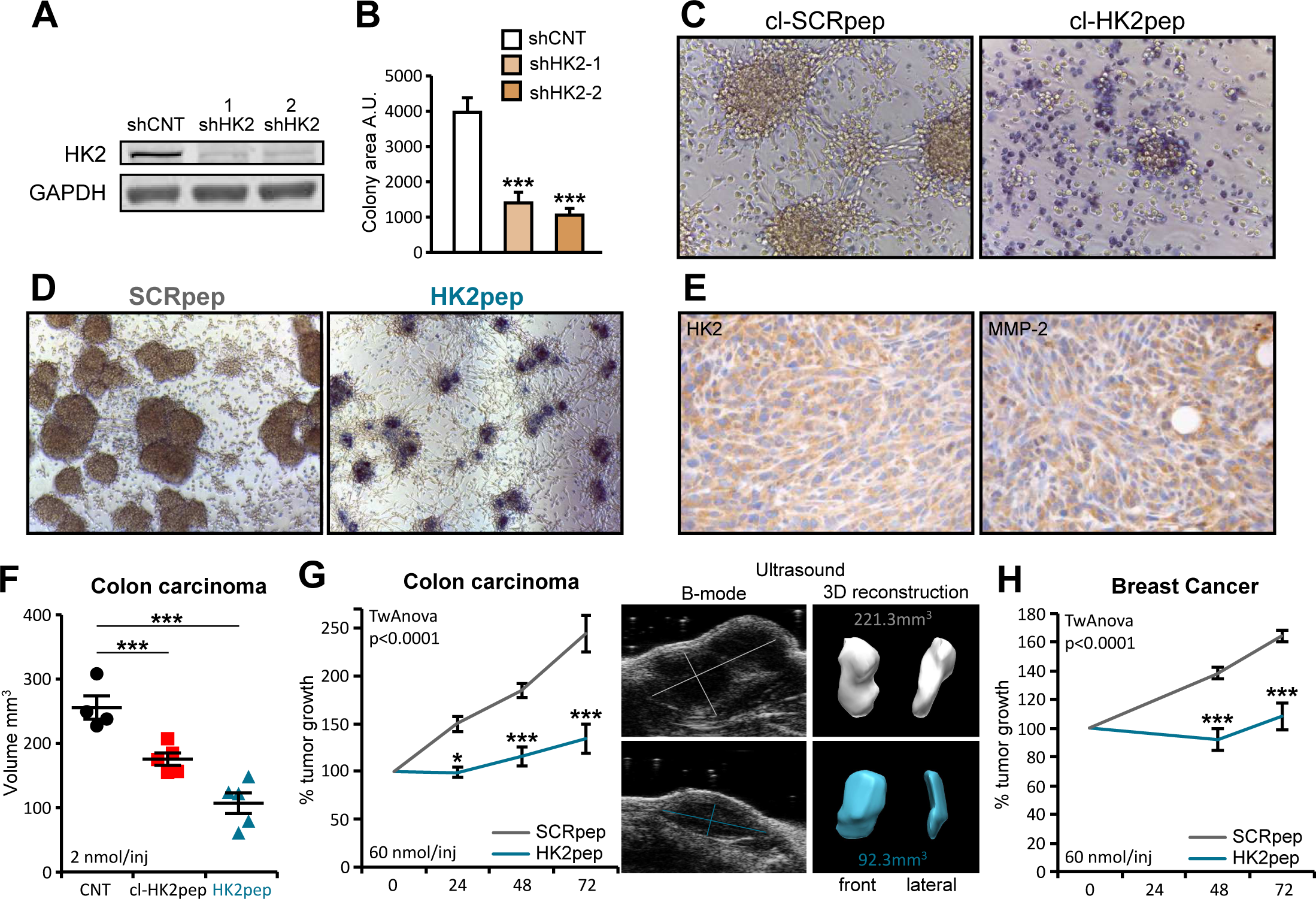
The HK2-targeting peptide inhibits *in vitro* and *in vivo* tumor growth. A-D Effect of HK2 targeting on *in vitro* tumorigenicity. HK2-silencing with two different shRNAs (shHK2-1 and shHK2-2) in CT26 colon cancer cells (A) inhibits growth in soft agar (B; colony area±SEM; n=6; Student’s t test ***p<0.001; A.U., arbitrary units). Treatment with cl-HK2pep (50 µM, 6 h; C) or with the uncleaved HK2pep (2 µM, 48 h; D) reverts formation of foci; dead cells are highlighted by Trypan Blue staining. E-H Effect of HK2-targeting on *in vivo* neoplastic growth. HK2- and MMP-2-expressing CT26 cells (E) are allografted in Balb/c male mice. Intratumor administration (5 injections of 2 nmol peptide every 12 h) of either cl-HK2pep or HK2pep (F), or intraperitoneal administration of HK2pep (G; 5 injections of 60 nmol peptide every 12 h) reduce neoplastic growth; the same effect is observed on allografts of breast carcinoma 4T1 cells (H; 5 injections of 60 nmol peptide every 12 h) in Balb/c female mice. Throughout the Figure, cl-SCRpep or SCRpep are used as peptide negative control. In G, representative ultrasound inspections and 3D reconstructions of tumor masses are shown (light grey: SCRpep treated animal; light blue: HK2pep treated animal). In F, mean tumor volume±SD is reported; Student’s t test ***p<0.001; in G, H, a two-way ANOVA is performed; Bonferroni post-test in the graphs *p<0.05; **p<0.01; ***p<0.001.

## Discussion

Our results indicate that HK2 specifically locates in MAMs, and that unhinging HK2 from MAMs swiftly elicits calpain-dependent death in cancer cells by funneling Ca^2+^ into mitochondria from ER and extracellular milieu. This signalling pathway is unprecedented, even though it is consistent with emerging evidences of a role played by MAMs in the maintenance of glucose homeostasis (Rieusset, 2018). We unveil a role of HK2 in handling intracellular Ca^2+^ fluxes, which suggests that HK2 can act as a regulatory hub to integrate glucose metabolism with Ca^2+^–mediated biochemical events, spanning from physiological responses to metabolic fluctuations to death induction under stress conditions. Hence, HK2 could contribute to the orchestration of the biochemical crosstalk between ER and mitochondria, with relevant implications both in the physiology and in several pathological conditions of HK2-expressing tissues, including diabetes, ischemic damage of the heart or muscle wasting diseases, notwithstanding tumors.

HK2 shuttling to mitochondria is modulated by Akt-dependent threonine phosphorylation (Miyamoto et al., 2008) and HectH9-dependent, non-proteolytic lysine ubiquitination (Lee *et al*, 2019). These data place mitochondrial localization of HK2 in a multifaceted network of regulatory events, and the functional interaction between HK2 and IP3Rs that we observe further adds to this network. A thorough dissection of the dynamic modulation of these interactions will be instrumental for the comprehension of the effects of HK2 displacement from MAMs. Importantly, non-tumor cells expressing HK2 are much more refractory to peptide toxicity than neoplastic cells. This is a promising observation under a therapeutic perspective. Nonetheless, it is necessary to understand whether MAM composition differs between tumor and non-tumor cells, and how this affects HK2 activity and localization, as well as to connect cell response to HK2pep to the pattern of calpain expression, in order to draw an exhaustive picture of the mode of peptide action and to preview its effectiveness in specific settings.

The use of CPPs as vectors for the delivery of molecules with anticancer activity constitutes an active field of investigation, even though the transfer of this approach to the clinical practice has been hampered up to now by several shortcomings, such as the lack of cell specificity and the short duration of action (Raucher & Ryu, 2015). Here we combine several strategies to overcome these limitations. HK2pep is an activatable peptide composed by assembled modular units that can be adapted to the specific tumor(s) to be targeted. The MMP2/9 cleavage site allows the selective activation of HK2pep where these metalloproteases are highly expressed, as in the microenvironment of several tumors, but cleavage sites for other extracellular matrix components can be envisaged and tailored to specific neoplastic types. The limited size of HK2 pep (less than 5 kDa) allows its passage through the fenestrated endothelium that characterizes many malignancies, and the rapidity of action of its cleaved form can circumvent problems connected with timing of degradation or excretion. Moreover, the absence of any inhibitory effect of HK2pep on the enzymatic activity of hexokinases minimizes the risk of non-specific damage caused by alterations of glucose metabolism at secondary sites. Hence, our approach to displace HK2 from MAMs integrates selectivity, efficacy and lack of off-target effects and could be evolved towards the development of novel anti-neoplastic strategies, conceivably in combination with other chemotherapeutic approaches that target different liabilities of neoplastic cells.

## Materials and Methods

### Cell culture and HK2 silencing

Human cervix carcinoma HeLa cells, human breast cancer MDA-MB-231 cells, human malignant peripheral nerve sheath tumor S462 cells, human plexiform neurofibroma PN 04.4 cells, mouse breast cancer 4T1 cells and mouse colorectal carcinoma CT26 cells, mouse macrophage RAW 264.7 cells and mouse myoblast C2C12 cells were cultured in DMEM medium (Gibco); colorectal carcinoma COLO 741 cells and B-CLL MEC1 cells were cultured in RPMI 1640 medium (Gibco); both culture media were supplemented with 10% Fetal Bovine Serum (Gibco), glutamine 2 mM (Gibco), penicillin-streptomycin (100 μg/ml; Invitrogen), and were kept at 37°C in a 5% CO_2_ humidified atmosphere.

HKII expression was stably interfered by infecting cells with a lentivirus carrying the following shRNAs (Sigma) against mouse HKII mRNA:

1. *CCGGCATCACCCTGCTGGTTCTAAACTCGAGTTTAGAACCAGCAGGGTGATGTTTTTG;*
2. *CCGGCGGTACAGAGAAAGGAGACTTCTCGAGAAGTCTCCTTTCTCTGTACCGTTTTTG. Scrambled shRNA (Sigma) was used as negative control. Transduced cells were selected in 1.0 µg/ml puromycin.*

### Histological and immunohistochemical analyses

Histological and immunohistochemical analyses were performed both on primary human samples (colorectal carcinoma, n=5; breast carcinoma, n=5; MPNST, n=5; lymph nodes deriving from B-cell chronic lymphocytic leukemia patients, n=5) and on samples derived from mouse tumor models (PN tumors; CT26 and 4T1 subcutaneous grafts in Balb/c mice). In detail, 4 µm-thick tissue sections were obtained from formalin-fixed paraffin-embedded tissue samples and representative tumor areas were selected on H&E-stained slides for immunohistochemical (IHC) analysis. IHC was performed using primary goat polyclonal anti-HK2 (sc-6521, Santa Cruz) and rabbit monoclonal anti-MPP2 (ab92536, Abcam) antibodies. Antigen retrieval was performed with heat/EDTA in the Bond-Max automated immunostainer (Leica Biosystems), as previously described (Pizzi *et al*, 2017).

### Cell lysates and Western immunoblot analysis

Cells lysis (5×10^5^ for each assay) was performed in Tris 20 mmol/l, NaCl 150 mmol/l, EDTA 2 mmol/l, EGTA 2 mmol/l, Triton X-100 0.5% supplemented with phosphatase and protease inhibitors (Sigma). Protein quantification was performed with BCA Protein Assay Kit (Thermo Scientific-Pierce). After SDS/PAGE gel electrophoresis, proteins were transferred onto nitrocellulose Hybond-C Extra membranes (Amersham, Uppsala; Sweden) and immunostained with goat polyclonal anti-HK2 (clone sc-6521, Santa Cruz), anti-GAPDH (#2118, Cell Signaling), ATP5A (ab14748, Abcam), anti-UQCRC2 (ab14745, Abcam), rabbit monoclonal anti-MMP2 (ab92536, Abcam), rabbit monoclonal anti-MMP9 (ab137867, Abcam) rabbit polyclonal and TOM20 (sc-11415, Santa Cruz) antibodies.

### Peptide synthesis

HK2pep, SCRpep, cl-HK2pep and cl-SCRpep were synthesized by automatic solid phase procedures using a multiple peptide synthesizer (SyroII, MultiSynTech GmbH) on a pre-loaded Wang resin (100-200 mesh) (Novabiochem). The fluoren-9-ylmethoxycarbonyl (Fmoc) strategy was used throughout the peptide chain assembly, utilizing O-(7-azabenzotriazol-1-yl)-N,N,N′,N′-tetramethyluronium hexafluorophosphate (HATU) as coupling reagent. The side-chain protected amino acid building blocks used were: Fmoc-Tyr(tert-butyl), Fmoc-Glu(tert-butyl), Fmoc-Ser(tertbutyl), Fmoc-Thr(tert-butyl), Fmoc-His(trityl), Fmoc-Asn(trityl), and Fmoc-Arg(2,2,4,6,7-pentamethyldihydrobenzofuran-5-sulfonyl). Cleavage of the peptides was performed by reacting the peptidyl-resins with a mixture containing TFA/H20/thioanisole/ethanedithiol/phenol (10 ml/0.5 ml/0.5 ml/0.25 ml/750 mg) for 2.5 h. Crude peptides were purified by a preparative reverse phase HPLC. Molecular masses of the peptide were confirmed by mass spectroscopy on a MALDI TOF-TOF using a Applied Biosystems 4800 mass spectrometer. The purity of the peptides was about 95% as evaluated by analytical reverse phase HPLC. Peptide sequences:

HK2pep: MIASHLLAYFFTELN-βA-RRRRRRRRR-PLGLAG-Ahx-EEEEEEEE

SCRpep: VGAHAGEYGAEALER-βA-RRRRRRRRR-PLGLAG-Ahx-EEEEEEEE

cl-HK2pep: MIASHLLAYFFTELN-βA-RRRRRRRRR-PLG

cl-SCRpep: VGAHAGEYGAEALER-βA-RRRRRRRRR-PLG

### Fluorescent peptide synthesis

Fluo-cl-HK2pep and fluo-cl-SCRpep were synthesized by microwave-assisted solid phase procedures using a Biotage Alstra peptide synthesizer on a pre-loaded HMPB-resin (100-200 mesh - Iris Biotech GmbH). The fluoren-9-ylmethoxycarbonyl (Fmoc) strategy was used throughout the peptide chain assembly. Activation of entering Fmoc-protected amino acids (0.3 M solution in DMF) was performed using 0.5 M Oxyma in DMF / 0.5 M DIC in DMF (1:1:1 molar ratio), with a 5 equivalent excess over the initial resin loading. Coupling steps were performed for 7 min at 75°C. Fmoc-deprotection steps were performed by treatment with a 20% piperidine solution in DMF at room temperature (1×10 min). Following each coupling or deprotection step, peptidyl-resin was washed with DMF (4×3.5ml). Upon complete chain assembly, resin was washed with DCM (5x 3.5ml) and gently dried under a nitrogen flow. Resin-bound peptide was treated with an ice-cold TFA, TIS, water, thioanisole mixture (90:5:2.5:2.5 v/v/v/v, 4 ml) for 2 hours. Crude peptides were purified by a preparative reverse phase HPLC. Molecular masses of the peptide were confirmed by mass spectroscopy using a Bruker Esquire 3000+ instrument equipped with an electro-spray ionization source and a quadrupole ion trap detector (QITD). The purity of the peptides was about 95% as evaluated by analytical reverse phase HPLC. Peptide conjugation with fluorescent dyes (ATTO) was performed by reacting peptide thiol groups with maleimide-ATTO dyes. Briefly, peptides were dissolved at a 5 mg/ml concentration in PBS buffer (pH 7.2) and 1.1 equivalents of maleimide-dye were added to the solution. Reaction was monitored by HPLC until completion, then the product isolated by preparative HPLC.

### HK2pep cleavage assay

Human recombinant MMP9 (911MP, R&D) was reconstituted following manufacturer’s instructions in a buffer composed by 50 mM Tris, 150 mM NaCl, 10 mM CaCl_2_, 0.05% Brij-35 (w/v) at pH 7.5 and activated with *p*-aminophenylmercuric acetate (Sigma). hMMP9 (20nM) was incubated for 4 hours with 1µg HK2pep in a final 100µl volume; samples were analyzed by mass/spec as described above.

### Measurement of hexokinase enzymatic activity

Hexokinase enzymatic activity was measured monitoring NADPH formation at 37°C with Infinite M200 spectrophotometer (TECAN) in a buffer containing 50mM Tris, 10mM MgCl_2_, 4mM ATP, 2mM glucose, 0.1 U/mL G6PDH and 1mM NADP, pH 7.4. Experiments were carried out either using Human HK2 recombinant protein (0.5µg; HXK0703, ATGEN) or total cell lysate (20µg) were exposed for 30 min to 10 μM cl-HK2pep. To discriminate between HK1 and HK2 activity in cells, samples were heated to 46°C for 30 min in order to inactivate thermo-sensitive HK2 activity.

### Immunofluorescence (IF) analyses

IF experiments were carried out 24 hours after transfection with TransIT®-LT1 (Mirus) on cells fixed in 4% PFA, quenched with NH_4_Cl (0.24% in PBS) and permeabilized with 0.1% Triton X-100. Primary antibodies (rabbit monoclonal anti-HK2, H.738.7, Thermo Scientific; mouse monoclonal anti-TOM20, sc-17764, Santa Cruz) were diluted 1:150 in blocking solution (2% BSA, 10% goat serum and 0.2% gelatin in PBS) and incubated for 90 min, at RT; AlexaFluor488/555/647-conjugated secondary antibodies (Life Technologies) were diluted 1:300 in blocking solution and incubated 45 min at RT. Images were collected on a Leica SP5-II confocal microscope, equipped with a 100x/1.4 N.A. Plan Apochromat objective. A WLL laser was used to excite each specific dye and a HyD (Leica) was employed for signal collection. After background-subtraction, the Pearson’s co-localization coefficient was calculated with the *ImageJ Co-localization* Analysis plugin. Displayed images were processed with the automatic ImageJ plugin *Enhance image* to improve signal intensity, but analyses were performed on raw, background-subtracted data. For measurements of co-localization with the split-GFP-based probe for ER-mitochondria contacts (SPLICS_L_) (Cieri et al., 2018), TOM20 and HK2 channels were background subtracted, a threshold (the mean fluorescence intensity for each cell) was imposed and the % of pixels co-localized with SPLICS_L_ was calculated, as described (Filadi *et al*, 2018). The plasmids encoding mt-YFP, mt-RFP and ER-CFP were previously described (Filadi et al., 2018).

### Mitochondrial Ca^2+^ measurements

For FRET-based ER Ca^2+^ measurements, cells were transfected with the ER-targeted D4ER Ca^2+^ probe and mounted into an open-topped chamber in Krebs–Ringer modified buffer (mKRB, in mM: 140 NaCl, 2.8 KCl, 2 MgCl_2_, 10 HEPES, 11 glucose, pH 7.4 at 37°C) supplemented with CaCl_2_ 1mM; imaging was performed with a DM6000 inverted microscope (Leica, HCX Plan Apo 40× oil objective, NA 1.25), as previously described (Fedeli *et al*, 2019). Excitation light was generated every 5s by a 410nm LED (Led Engin LZ1-00UA00 LED). ImageJ was used for off-line analysis of FRET experiments. YFP and CFP images were background-subtracted and analyzed selecting specific regions of interest (ROIs) on each cell. The ratio (R) between YFP and CFP emissions was calculated for each frame.

For GCAMP6f Ca^2+^ measurements, cells were transfected with a cDNA encoding mitochondrial and nuclear GCAMP6f (Filadi et al., 2018). To perform Ca^2+^ measurements, medium was replaced with mKRB buffer supplemented with 1µM CsH (Adipogen) and with 1mM CaCl_2_ or 500µM EGTA. Where indicated, specific drugs were added to the buffer and pre-incubated for 30 min with BAPTA-AM (5µM, Thermo Fisher Scientific); or 1 hour with Xestospongin C (5µM, Santa Cruz), CsA (5µM, Sigma), C63 (5µM) or PD150606 (50µM, Calbiochem). Fluorescence was recorded with an inverted microscope (Zeiss Axiovert 100, Fluar 40× oil objective, NA 1.30) in the 500-530nm range (by a band-pass filter, Chroma Technologies). Probes were sequentially excited at 475 nm and at 410 nm, respectively for 180 and 300ms, every 5s. Excitation light, produced by a monochromator (polychrome V; TILL Photonics), was filtered with a 505 nm DRLP filter (Chroma Technologies). After background subtraction, images were analyzed with ImageJ, calculating the ratio (R) between emissions generated by exciting cells at 475 and 410 nm, respectively, in specific ROIs comprising the entire mitochondrial network. [Ca^2+^] is proportional to R.

For aequorin Ca^2+^ measurements, cells were transfected with a plasmid encoding low-affinity mitochondrial matrix aequorin (mit-Aeqmut), which was reconstituted by incubating cells for 1h in mKRB, supplemented with CaCl_2_ 1 mM and native coelenterazine (5µM, Biotium). Measurements were performed as previously described (Filadi et al., 2018).

### IP_3_ measurement assay

In order to assess changes in intracellular IP3 levels, cells were transfected with a plasmid encoding the GFP-tagged pleckstrin homology (PH) domain of PLC-δ1 (GFP-PHD). Quantification of IP_3_ generation was evaluated as variations in cytosolic GFP-PHD fluorescence (F) normalized to initial cytosolic fluorescence (F0), due to the release of GFP-PHD from the plasma membrane upon PIP2 cleavage and IP3 generation (Hirose *et al*, 1999). Only cells (∼ 80%) in which a plasma membrane localization of GFP-PHD was evident were selected for analysis. Images were collected on a DM6000 inverted microscope (Leica, HCX Plan Apo 40× oil objective, NA 1.25). Excitation light was generated every 5 s by a 460 nm LED (Led Engin).

### Measurements of mitochondrial membrane potential

Mitochondrial membrane potential was analyzed using the fluorescent potentiometric compound tetramethylrhodamine methyl ester (TMRM, 20nM; Invitrogen). Cells were incubated in mKRB or DMEM without phenol red and with CsH (1μM) to inhibit P-glycoproteins. Recordings were performed either with a fluorescence microscope (Olympus IX71 or Leica DMI600B) or with a flow cytometer (FACSCanto II, Becton Dickinson). For live microscope experiments, recordings were carried out every 30 sec using CellR software (Olympus) or LAS AF software (Leica). For flow cytometry analyses, TMRM signals are measured every 15 min and analyzed with FACSDiva software (Becton Dickinson). Movies were realized using CellR software (Olympus).

### Cell viability assays

Cell viability was assessed either by cytofluorimetry or by fluorescence microscopy. Cytofluorimetric recordings of phosphatidylserine exposure on the cell surface (increased staining of FITC-conjugated Annexin-V; Roche) and loss of plasma membrane integrity (7-AAD staining; Sigma) were repeated every 15 min on a FACSCanto II instrument and analyzed with FACSDiva software. Double negative cells were considered viable. Fluorescence microscope recordings were performed with Leica DMI600B microscope and acquired with LAS AF software in the presence of Annexin V-FITC, 7-AAD and TMRM pre-incubated for 45 min. Images and movie obtained with LAS AF software were analyzed with ImageJ software.

### Isolation of B lymphocytes

Cell samples that matched standard morphological and immunophenotypic criteria for B-CLL were collected from 22 therapy-free patients after obtaining informed consent according to the Declaration of Helsinki. Patient characteristics are reported in Table 1. Cells were separated by Ficoll-Hypaque gradient centrifugation (Euroclone) and B cells were purified from the PBMCs by removing T cells with the sheep erythrocyte rosetting method (Frezzato *et al*, 2014). Untreated peripheral blood B cells were isolated from the PBMCs of healthy donors by negative selection with the RosetteSep for isolation of B cells (StemCell Technologies). All samples were used when they contained ≥95% CD19^+^ cells, as assessed by flow cytometry.

### *In vitro* tumorigenesis assays

For soft agar assays, cells in DMEM medium supplemented with 0.5% serum were mixed with low melting point agarose (Promega) at a final 0.6% concentration and plated on a bottom layer of DMEM with 1% LMP agarose. Fresh medium (DMEM 4% serum) was added every 3^rd^ day. At day 15^th^ colonies were stained with Crystal Violet 0.005% and analyzed with ImageJ software.

For focus forming assays, cells were seeded in standard DMEM medium with 10% FBS, which was replaced with DMEM 0.5% FBS after cells reached confluence. Treatment with peptides were performed after formation of foci; dead cells were highlighted by Trypan Blue staining. Images were acquired using a Leica DMIL LED microscope equipped with a Leica ICC50 HD camera.

### *In vivo* tumorigenesis assays

Colon cancer CT26 cells or breast cancer 4T1 cells were allografted in Balb/c mice according to a protocol approved from Italian Health Ministry (authorization number: 547/2016-PR). Briefly, 100.000 CT26 cells or 200.000 4T1 cells were injected subcutaneously in the flank of male or female animals, respectively, in a 100 μl final volume containing 30% Matrigel (Corning). Tumors were visible under the skin after 7-9 days and measured with caliper (two major axes). Tumor volume was calculated using the formula: (a*b^2^)/2. Ultrasound inspections of representative tumors were performed using Vevo 2100 (Visualsonic, Fujifilm). Acquisitions, measurements and 3D-reconstructions of tumors were done using VevoLAB 1.7.0.7071 software. Peptides where injected as described in the text.

## Acknowledgements

This work was supported by grants from University of Padova, Neurofibromatosis Therapeutic Acceleration Program, Associazione Italiana Ricerca Cancro (AIRC grant IG 2017/20749 to A.R. and AIRC grant IG 2015/17067 to P.B.), Piano for Life Onlus and Linfa OdV. IM was recipient of a Young Investigator Award Grant from the Children’s Tumor Foundation.

## Author Contributions

Conceptualization, F.C., A.R. and P.B.; Investigation, F.C., R.F., I.M and M.P.; Resources, O.M., N.D., A.G., F.F. and L.T.; Formal Analysis, F.C. and R.F.; Visualization, F.C. and R.F.; Funding Acquisition, A.R. and P.B.; Supervision, F.C. and A.R.; Project Administration, A.R.; Writing-Original Draft, A.R.; Writing-Review and Editing, A.R., F.C., R.F., P.P., L.T. and P.B.

## Expanded view figure legends

**Figure EV1. Related to Figure 1. HK2 locates in MAMs of cancer cells.**

A, B Fluorescence co-staining of HK2 and split-GFP-based probe for ER-mitochondria contacts (SPLICS_L_) on colorectal carcinoma COLO 741 cells, breast cancer MDA-MB-231 cells, plexiform neurofibroma PN 04.4 cells and malignant peripheral nerve sheath tumor S462 cells (A) and on HeLa cells, where the outer mitochondrial membrane marker TOM20 is also analyzed for its co-localization with MAMs (B). HK2 is revealed with an AlexaFluor555-conjugated secondary antibody, TOM20 with an AlexaFluor647-conjugated secondary antibody; the merged signal is white in both analyses. MAM localization of HK2 or TOM20 is quantified in the bar graph on the right. Scale bar: 15µM.

**Figure EV2. Related to Figure 1. Characterization of cl-HK2pep.**

A HeLa cells expressing mitochondrial-targeted RFP are treated for 2 min with either cl-SCRpep or cl-HK2pep (1 μM each) labeled with the green fluorophore ATTO 488. The yellow signal indicates mitochondrial localization of the peptide. Scale bar: 15 µM.

B HK2 displacement from HeLa mitochondria after a 2 min treatment with cl-HK2pep is shown by loss of merging signal analyzed as in (Figure 1C). cl-SCRpep is used as a negative control and Pearson’s co-localization coefficient is indicated in figure (cl-SCRpep n=40 cells; cl-HK2pep n=51 cells; p<0.01 with a Student’s *t* test). Scale bar: 15µM.

C Effect of 10 μM cl-HK2pep on glucose phosphorylation in CT26 cell extracts, where both total hexokinase activity and HK1/HK2 specific activities are measured (n=3).

**Figure EV3. Related to Figure 2. Kinetics of cl-HK2pep-induced mitochondrial depolarization.**

A IP_3_ levels are assessed with the GFP-PHD probe in presence or not of 10 μM BAPTA-AM; data are reported as mean of fluorescent signals±SEM (n=3)

B-E Changes in mitochondrial membrane potential following cl-HK2pep treatment assessed with TMRM as in Figure 2G. In (B, C) HeLa cells are treated with 5 μM cl-HK2pep; mitochondrial depolarization is shown for single cells (B; n=60) or as an averaged trace (C; n=60). TMRM signal is normalized to the initial value and expressed in percentage for each time point. D, TMRM positive HeLa cells are expressed in percentage (threshold count: fluorescence signal >40% of initial value; n=60). E, HeLa cells are treated with 2 μM cl-HK2pep (n=216) and analyzed as in (C).

F, G Changes in Ca^2+^ levels and membrane potential in HeLa cell mitochondria following treatment with 2 µM cl-HK2pep. Measurements are performed in parallel on cells expressing mito-GCAMP6f and loaded with TMRM. Two different single cell traces are shown, characterized by fast (F) or slow (G) mitochondrial depolarization following peptide treatment.

H Mitochondrial depolarization in 4T1 cells treated with 2 μM cl-HK2pep (n=129) is analyzed as in (C). Where indicated, BAPTA-AM (10 µM; n=147) or Xe-C (5 µM; n=141) are added as in Figure 2B 1 hour before experiment.

**Figure EV4. Related to Figure 3. Cell death induction by cl-HK2pep in different cancer cell models**.

A Western immunoblot analysis of HK2 levels in murine cell models (colon carcinoma CT26 cells, breast cancer 4T1 cells, macrophage RAW 264.7 cells and myoblast C2C12 cells). ATP5A and GAPDH are mitochondrial and cytosolic loading controls, respectively.

B, C Effect of cl-HK2pep (2 and 5 μM) on cell death in RAW 264.7 and C2C12 non transformed cells. Cell death is assessed by cytofluorimetry as in Figure 3C.

D-F The effect of the pan-caspase inhibitor Z-VAD-fmk (50 µM) on death elicited in HeLa, CT26 and 4T1 cells by a 15 min treatment with 5 µM cl-HK2pep is assessed as in B.

G, H The effect of the pan-calpain inhibitor PD150606 (50 μM) on mitochondrial depolarization elicited by 5µM cl-HK2pep is measured as in Figure 3B (n=3 in each cell line; mean percentage of dead cells is normalized to time 0±SD; Two-way ANOVA: p<0.0001 for both examined cell lines; Bonferroni post-test in graphs ***p<0.001).

I-M Effect of PD150606 on cell death elicited by treatment with 5 µM cl-HK2pep in CT26, 4T1, COLO 741, MDA-MB-231 and MEC1 cancer cell models is analyzed as in (B). In (G-M), data ±SD are analyzed with a Two-way ANOVA: p<0.0001 for all examined cell lines; Bonferroni post-test in graphs ***p<0.001. In experiments throughout the Figure, cl-SCRpep is used as a negative control; n≥3 for each cell line.

**Figure EV5. Related to Figure 5. Analysis of mouse samples following peptide treatment.**

A Western immunoblot analysis of MMP2 and MMP9 expression in neoplasms formed by 4T1 cells in mouse allografts; TOM20 is used as loading control.

B Representative Hematoxylin/Eosin staining on samples from heart, skeletal muscle, kidney and spleen obtained from mice bearing CT26 tumors after peptide treatment as in Figure 4M.

## Movie legends

**Movie 1. Related to Figure 2. cl-HK2pep induces Ca**^**2+**^ **rise in mitochondria.**

HeLa cells expressing the mitochondrial-GCAMP6f Ca^2+^ sensor are treated with 2 μM cl-HK2pep at minute 2; fluorescence is recorded every 5 sec.

**Movie 2. Related to Figure 2. Effect of cl-HK2pep (5 μM) treatment on mitochondrial membrane potential.**

Mitochondrial membrane potential of HeLa cells assessed with the TMRM probe; fluorescence is recorder every 30 sec. Cells are treated with cl-HK2pep 5 μM at minute 4.

**Movie 3. Related to Figure 2. Effect of cl-HK2pep (2 μM) treatment on mitochondrial membrane potential.**

Mitochondrial membrane potential of HeLa cells assessed with the TMRM probe; fluorescence is recorder every 30 sec. Cells are treated with cl-HK2pep 2 μM at minute 4.

**Movie 4. Related to Figure 2. Effect of cl-HK2pep and XeC treatment on mitochondrial membrane potential.**

Mitochondrial membrane potential of HeLa cells assessed with the TMRM probe; fluorescence is recorder every 30 sec. Cells are preincubated 1h with Xe-C (5 μM) and treated with cl-HK2pep 2 μM at minute 4.

**Movie 5. Related to Figure 3. Effect of cl-HK2pep and CsA treatment on mitochondrial membrane potential.**

Mitochondrial membrane potential of HeLa cells assessed with the TMRM probe; fluorescence is recorder every 30 sec. Cells are preincubated 1h with CsA (5 μM) and treated with cl-HK2pep 2 μM at minute 4.

**Movie 6. Related to Figure 3. Effect of cl-HK2pep and C63 treatment on mitochondrial membrane potential.**

Mitochondrial membrane potential of HeLa cells assessed with the TMRM probe; fluorescence is recorder every 30 sec. Cells are preincubated 1h with C63 (5 μM) and treated with cl-HK2pep 2 μM at minute 4

**Movie 7. Related to Figure 3. Kinetic analysis of cell death induced by cl-HK2pep.**

HeLa cells are loaded with TMRM and kept in a medium containing Annexin V-FITC and 7-AAD. cl-HK2pep (5 μM) is added at minute 2 and fluorescence is recorded every 2 sec.

**Movie 8. Related to Figure 5. Representative 3D tumor reconstructions (control conditions).** Tumors induced by allograft injection of CT26 cells are ultrasound inspected and scanned using an automated motor for 3D recordings. Tumors are treated with SCRpep as in Fig. 5G.

**Movie 9. Related to Figure 5. Representative 3D tumor reconstructions (HK2pep treatment).** Tumors induced by allograft injection of CT26 cells are ultrasound inspected and scanned using an automated motor for 3D recordings. Tumors are treated with HK2pep as in Fig. 5G.

## References

Akins NS, Nielson TC, Le HV (2018) Inhibition of Glycolysis and Glutaminolysis: An Emerging Drug Discovery Approach to Combat Cancer. Curr Top Med Chem 18: 494–504

Bernardi P, Rasola A, Forte M, Lippe G (2015) The Mitochondrial Permeability Transition Pore: Channel Formation by F-ATP Synthase, Integration in Signal Transduction, and Role in Pathophysiology. Physiol Rev 95: 1111–1155

Berridge MJ, Bootman MD, Roderick HL (2003) Calcium signalling: dynamics, homeostasis and remodelling. Nat Rev Mol Cell Biol 4: 517–529

Bhalla K, Jaber S, Nahid MN, Underwood K, Beheshti A, Landon A, Bhandary B, Bastian P, Evens AM, Haley J, Polster B, Gartenhaus RB (2018) Role of hypoxia in Diffuse Large B-cell Lymphoma: Metabolic repression and selective translation of HK2 facilitates development of DLBCL. Sci Rep 8: 744

Cannino G, Ciscato F, Masgras I, Sanchez-Martin C, Rasola A (2018) Metabolic Plasticity of Tumor Cell Mitochondria. Front Oncol 8: 333

Chiara F, Castellaro D, Marin O, Petronilli V, Brusilow WS, Juhaszova M, Sollott SJ, Forte M, Bernardi P, Rasola A (2008) Hexokinase II detachment from mitochondria triggers apoptosis through the permeability transition pore independent of voltage-dependent anion channels. PLoS ONE 3: e1852

Cieri D, Vicario M, Giacomello M, Vallese F, Filadi R, Wagner T, Pozzan T, Pizzo P, Scorrano L, Brini M, Cali T (2018) SPLICS: a split green fluorescent protein-based contact site sensor for narrow and wide heterotypic organelle juxtaposition. Cell Death Differ 25: 1131–1145

Csordas G, Weaver D, Hajnoczky G (2018) Endoplasmic Reticulum-Mitochondrial Contactology: Structure and Signaling Functions. Trends Cell Biol 28: 523–540

DeWaal D, Nogueira V, Terry AR, Patra KC, Jeon SM, Guzman G, Au J, Long CP, Antoniewicz MR, Hay N (2018) Hexokinase-2 depletion inhibits glycolysis and induces oxidative phosphorylation in hepatocellular carcinoma and sensitizes to metformin. Nat Commun 9: 446

Fabbri G, Dalla-Favera R (2016) The molecular pathogenesis of chronic lymphocytic leukaemia. Nat Rev Cancer 16: 145–162

Fedeli C, Filadi R, Rossi A, Mammucari C, Pizzo P (2019) PSEN2 (presenilin 2) mutants linked to familial Alzheimer disease impair autophagy by altering Ca(2+) homeostasis. Autophagy: 1–19

Filadi R, Leal NS, Schreiner B, Rossi A, Dentoni G, Pinho CM, Wiehager B, Cieri D, Cali T, Pizzo P, Ankarcrona M (2018) TOM70 Sustains Cell Bioenergetics by Promoting IP3R3-Mediated ER to Mitochondria Ca(2+) Transfer. Curr Biol 28: 369–382 e366

Filadi R, Theurey P, Pizzo P (2017) The endoplasmic reticulum-mitochondria coupling in health and disease: Molecules, functions and significance. Cell Calcium 62: 1–15

Frezzato F, Trimarco V, Martini V, Gattazzo C, Ave E, Visentin A, Cabrelle A, Olivieri V, Zambello R, Facco M, Zonta F, Cristiani A, Brunati AM, Moro S, Semenzato G, Trentin L (2014) Leukaemic cells from chronic lymphocytic leukaemia patients undergo apoptosis following microtubule depolymerization and Lyn inhibition by nocodazole. Br J Haematol 165: 659–672

Gu JJ, Singh A, Xue K, Mavis C, Barth M, Yanamadala V, Lenz P, Grau M, Lenz G, Czuczman MS, Hernandez-Ilizaliturri FJ (2018) Up-regulation of hexokinase II contributes to rituximab-chemotherapy resistance and is a clinically relevant target for therapeutic development. Oncotarget 9: 4020–4033

Hirose K, Kadowaki S, Tanabe M, Takeshima H, Iino M (1999) Spatiotemporal dynamics of inositol 1,4,5-trisphosphate that underlies complex Ca2+ mobilization patterns. Science 284: 1527-1530

Isaacson KJ, Martin Jensen M, Subrahmanyam NB, Ghandehari H (2017) Matrix-metalloproteinases as targets for controlled delivery in cancer: An analysis of upregulation and expression. J Control Release 259: 62–75

Lee HJ, Li CF, Ruan D, He J, Montal ED, Lorenz S, Girnun GD, Chan CH (2019) Non-proteolytic ubiquitination of Hexokinase 2 by HectH9 controls tumor metabolism and cancer stem cell expansion. Nat Commun 10: 2625

Martini V, Gattazzo C, Frezzato F, Trimarco V, Pizzi M, Chiodin G, Severin F, Scomazzon E, Guzzardo V, Saraggi D, Raggi F, Martinello L, Facco M, Visentin A, Piazza F, Brunati AM, Semenzato G, Trentin L (2017) Cortactin, a Lyn substrate, is a checkpoint molecule at the intersection of BCR and CXCR4 signalling pathway in chronic lymphocytic leukaemia cells. Br J Haematol 178: 81–93

Masgras I, Rasola A, Bernardi P (2012) Induction of the permeability transition pore in cells depleted of mitochondrial DNA. Biochim Biophys Acta 1817: 1860–1866

Mathupala SP, Ko YH, Pedersen PL (2009) Hexokinase-2 bound to mitochondria: cancer’s stygian link to the “Warburg Effect” and a pivotal target for effective therapy. Semin Cancer Biol 19: 17–24

Mathupala SP, Ko YH, Pedersen PL (2010) The pivotal roles of mitochondria in cancer: Warburg and beyond and encouraging prospects for effective therapies. Biochim Biophys Acta 1797: 1225–1230

Miyamoto S, Murphy AN, Brown JH (2008) Akt mediates mitochondrial protection in cardiomyocytes through phosphorylation of mitochondrial hexokinase-II. Cell Death Differ 15: 521–529

Olson ES, Aguilera TA, Jiang T, Ellies LG, Nguyen QT, Wong EH, Gross LA, Tsien RY (2009) In vivo characterization of activatable cell penetrating peptides for targeting protease activity in cancer. Integr Biol (Camb) 1: 382–393

Pantic B, Trevisan E, Citta A, Rigobello MP, Marin O, Bernardi P, Salvatori S, Rasola A (2013) Myotonic dystrophy protein kinase (DMPK) prevents ROS-induced cell death by assembling a hexokinase II-Src complex on the mitochondrial surface. Cell Death Dis 4: e858

Patra KC, Wang Q, Bhaskar PT, Miller L, Wang Z, Wheaton W, Chandel N, Laakso M, Muller WJ, Allen EL, Jha AK, Smolen GA, Clasquin MF, Robey RB, Hay N (2013) Hexokinase 2 is required for tumor initiation and maintenance and its systemic deletion is therapeutic in mouse models of cancer. Cancer Cell 24: 213–228

Pizzi M, Agostinelli C, Righi S, Gazzola A, Mannu C, Galuppini F, Fassan M, Visentin A, Piazza F, Semenzato GC, Rugge M, Sabattini E (2017) Aberrant expression of CD10 and BCL6 in mantle cell lymphoma. Histopathology 71: 769–777

Prudent J, McBride HM (2017) The mitochondria-endoplasmic reticulum contact sites: a signalling platform for cell death. Curr Opin Cell Biol 47: 52–63

Rasola A, Bernardi P (2011) Mitochondrial permeability transition in Ca(2+)-dependent apoptosis and necrosis. Cell Calcium 50: 222–233

Rasola A, Bernardi P (2014) The mitochondrial permeability transition pore and its adaptive responses in tumor cells. Cell Calcium 56: 437–445

Raucher D, Ryu JS (2015) Cell-penetrating peptides: strategies for anticancer treatment. Trends Mol Med 21: 560–570

Rebecchi MJ, Pentyala SN (2000) Structure, function, and control of phosphoinositide-specific phospholipase C. Physiol Rev 80: 1291–1335

Rieusset J (2018) The role of endoplasmic reticulum-mitochondria contact sites in the control of glucose homeostasis: an update. Cell Death Dis 9: 388

Roberts DJ, Miyamoto S (2015) Hexokinase II integrates energy metabolism and cellular protection: Akting on mitochondria and TORCing to autophagy. Cell Death Differ 22: 248–257

Roberts DJ, Tan-Sah VP, Ding EY, Smith JM, Miyamoto S (2014) Hexokinase-II positively regulates glucose starvation-induced autophagy through TORC1 inhibition. Mol Cell 53: 521–533

Robey RB, Hay N (2006) Mitochondrial hexokinases, novel mediators of the antiapoptotic effects of growth factors and Akt. Oncogene 25: 4683–4696

Semenza GL (2013) HIF-1 mediates metabolic responses to intratumoral hypoxia and oncogenic mutations. J Clin Invest 123: 3664–3671

Storr SJ, Carragher NO, Frame MC, Parr T, Martin SG (2011) The calpain system and cancer. Nat Rev Cancer 11: 364–374

Suh DH, Kim MA, Kim H, Kim MK, Kim HS, Chung HH, Kim YB, Song YS (2014) Association of overexpression of hexokinase II with chemoresistance in epithelial ovarian cancer. Clin Exp Med 14: 345–353

Vander Heiden MG, DeBerardinis RJ (2017) Understanding the Intersections between Metabolism and Cancer Biology. Cell 168: 657–669

Wang L, Xiong H, Wu F, Zhang Y, Wang J, Zhao L, Guo X, Chang LJ, Zhang Y, You MJ, Koochekpour S, Saleem M, Huang H, Lu J, Deng Y (2014) Hexokinase 2-mediated Warburg effect is required for PTEN- and p53-deficiency-driven prostate cancer growth. Cell Rep 8: 1461–1474

Wilson JE (2003) Isozymes of mammalian hexokinase: structure, subcellular localization and metabolic function. J Exp Biol 206: 2049–2057

Wolf A, Agnihotri S, Micallef J, Mukherjee J, Sabha N, Cairns R, Hawkins C, Guha A (2011) Hexokinase 2 is a key mediator of aerobic glycolysis and promotes tumor growth in human glioblastoma multiforme. J Exp Med 208: 313–326

